# MAIS: an in-vitro sandbox enables adaptive neuromodulation via scalable neural interfaces

**DOI:** 10.1101/2025.03.15.641656

**Authors:** Haoman Chen, Fanxuan Chen, Xinyu Chen, Yang Liu, Junpeng Xu, Jiajun Li, Xueying Bao, Yuzhe Chen, Haojun Sun, Jiaju Jiang, Fangzhou Ye, Jianzhong Su, Gen Yang, Fangfu Ye, Zhouguang Wang, Liyu Liu, Saiyin Hexige, Xiaokun Li, Lixiang Ma, Jianwei Shuai

## Abstract

Brain-machine interfaces (BMIs) predominantly rely on static digital architectures to decode biological neuronal networks, a paradigm that is incompatible with natural neural coding in the human brain^1–4^. Bridging this gap is a critical step in combating neuronal dysfunction, enhancing brain functionality, and refining the precision of neuroprosthetics^5^. The integration of brain organoids with microelectrode array (MEA), as a class of BMIs, offers a humanized in vitro platform with unique biological compatibility advantages for dynamic neuronal decoding. This study resolves the biological-electronic encoding incompatibility of brain organoid-MEA Integration through three progressive breakthroughs. First, a human-machine hybrid agent is developed as a newly proposed bioengineered platform that couples brain organoids together with high-density MEAs and computational chips, enabling closed-loop perturbation of biological neuronal networks via exogenous signals. Second, through plasticity-driven real-time tracking of neuronal activity, we establish dynamically reconfigurable stimulation nodes that self-align with the electrophysiological states of the organoids. This resolves the exogenous-endogenous encoding mismatch by implementing plasticity-driven adaptation principles that ensure biological compatibility through spatially adaptive coordination. Finally, through shared plasticity rules rather than centralized control, we construct the first scalable multi-agent interactive system (MAIS) and demonstrate its real-world applications. Through designed scenarios of pathological/normal neuronal network interaction, we validate that MAIS achieves stable cross-network coordination. MAIS embodies a self-evolving neural coding sandbox in which plasticity-driven dynamic decoding bridges the compatibility gaps between biological and digital systems, providing a scalable and foundational infrastructure for human-centered neural interfaces.

## Main

Human-derived neural cultured organoid is an important component of the in vitro system, complemented by corresponding stimulatory feedback research frameworks, which are fundamental building blocks for in-vitro brain-machine interfaces (BMIs) ^6–10^. Recently, in vitro systems have gained prominence due to their ability to bridge insights from biology, artificial intelligence, and medicine, while addressing critical limitations of BMIs as research sandboxes^11^. The convergence of advances in the understanding of functional neuronal networks and advances in electrode technology has increased the accessibility of in vitro systems for exploring complex brain connections, which is essential for the development of adaptive BMI architectures^12–14^. In contrast to clinical studies, which are limited by technical and ethical challenges, in vitro systems provide opportunities to investigate the neurobiological basis of human-specific adaptations through controlled prototyping of BMIs^15–17^. In addition, their compatibility with high-throughput experimental platforms allows for in-depth exploration of neuronal dynamics and targeted modulation of neuronal dysfunction via BMIs-compatible intervention strategies, enabling research in diverse areas not readily achievable with traditional in vivo methods^18–21^. However, existing systems often rely on artificial spatial design during neuronal perturbation that is poorly suited to the self-organizing nature of biological networks, creating fundamental discrepancies between experimental paradigms and intrinsic neurodynamic principles —a critical barrier to unraveling complex neurological phenomena^13,22–24^.

Among various in vitro systems, the integration of brain organoids with microelectrode arrays (MEAs) has emerged as a key platform for merging carbon-based biological entities with silicon-based computational systems. Derived from pluripotent stem cells, brain organoids recapitulate critical features of the human brain, such as neuronal network formation, synaptic plasticity, and spontaneous electrical activity, which are critical for the development of BMIs^25–28^. When coupled with high-density MEA technology, these hybrid systems enable real-time recording and perturbation of neuronal circuits, providing unique insights into brain-like signal processing and neural computation through BMI-compatible interfaces^12,14^. Recent studies have highlighted the transformative potential of organoid-MEA platforms in various applications to advance BMI paradigms. For example, researchers have used this approach to achieve adaptive reservoir computing with brain organoids, demonstrating their ability to process spatiotemporal information and perform tasks such as speech recognition and nonlinear equation prediction through biohybrid BMIs implementations^22^. In addition, in vitro neuronal networks have shown that task-specific training, such as engaging neurons in the game “Pong”, can induce dynamic behaviors close to criticality, linking task performance to structured sensory input^23^. Another major advance is the integration of neuronal networks into simulated environments, where closed-loop feedback facilitates goal-directed learning and reveals the capacity for self-organization in neuronal cultures^24^. Collectively, these findings highlight the versatility and adaptability of organoid MEA systems for exploring the dynamic phenomena associated with neural interfaces and their emergent properties.

However, two fundamental limitations remain: First, existing platforms primarily operate in single-organoid paradigms, missing opportunities to study properties arising from inter-agent communication that may mimic complex interpersonal communication. Second, most stimulation protocols employ static spatial patterns that conflict with dynamic neuronal networks, highlighting the need for adaptive frameworks aligned with intrinsic neuroplasticity mechanisms to achieve dynamic symbiosis in BMIs. These two gaps precisely define the frontier for the development of next-generation BMI systems. Addressing these limitations requires fundamentally new approaches to establish programmable interfaces for spatiotemporal activity modulation and to create interconnected multi-organoid architectures manifesting complex group behaviors. This forms the conceptual basis for the construction of brain organoid-based multi-agent systems with multidisciplinary application potential.

In this study, we present the Multi-Agent Interactive System (MAIS)—a biologically aligned framework that addresses the two fundamental limitations of existing BMI platforms. By integrating human cortical organoids with reprogrammable microelectrode array chips and computing chips, MAIS establishes the first dynamic closed-loop perturbation system capable of both spatial programming control and multi-organoid emergent behavior simulation, redefining the scalability of BMIs. Within this architecture, a network of biohybrid agents-each consisting of hCOs functionally fused with MEA substrates and computational chips-engage in role-specific perturbative interactions that mirror highly human interpersonal communication. Using isogenic normal and disease model hCOs, including autism spectrum disorder and Huntington’s disease variants, MAIS implements three core agent types: 1) “Doctor Agents” that perform differential diagnosis through neural pattern recognition, 2) “Patient Agents” that manifest pathological phenotypes, and 3) “Health Agents” that maintain normative activity baselines, establishing the first scalable platform for studying network-level neurological dysfunction through targeted multi-agent perturbation (Fig. 1a). Bioelectronic coupling is provided by silicon substrates that translate biological neuronal signals into programmable stimulation patterns. Real-time neuronal dynamics captured by MEAs are processed by adaptive algorithms that generate context-specific electrical perturbations, implementing the reinforcement learning paradigm through neuroplasticity-oriented stimulation protocols (Fig. 1b). These neuroplasticity-aligned stimulation protocols enable MAIS to make the critical transition from artificial spatial constraints to biologically compatible closed-loop regulation, a key advancement highlighted in our spatial programming methodology. The perturbation framework employs a biologically grounded reinforcement learning architecture in which agent-environment interactions are modeled by dynamic state-action mappings, setting new standards for BMI-responsive interfaces, with neuronal activity patterns serving as both input states and modulable outputs (Fig. 1c). In multiagent configurations, the direct representation of neuronal activity through a shared electrophysiological features enable naturalistic interactions while maintaining individual agent specificity (Fig. 1d). Crucially, the “Doctor Agent” performs continuous differential analysis of the dynamics of the “Patient” and “Control” agents, and delivers plasticity-inducing stimulation patterns that recalibrate pathological activity toward normative baselines. Through iterative cycles of pattern comparison, adaptive stimulation, and neural plasticity, MAIS demonstrates the first implementation of self-organizing therapeutic modulation in vitro, establishing a engineering platform for investigating neuronal dysfunction, developing biohybrid intelligence systems, and prototyping next-generation neural interfaces.

**Figure 1.**
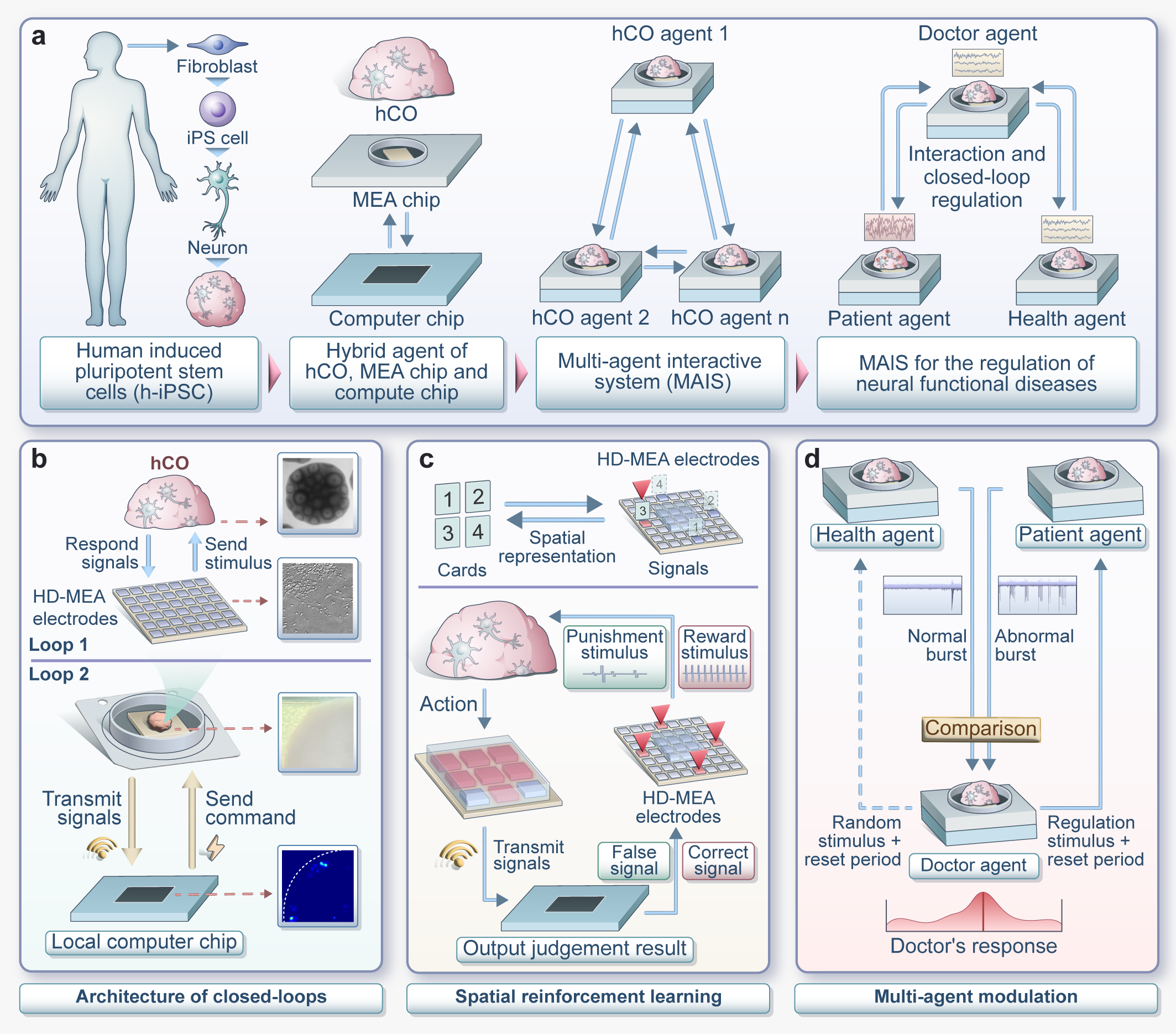
Conceptual framework and core architecture of the Multi-Agent Interactive System (MAIS) **a**) Developmental workflow of the Multi-Agent Interactive System (MAIS). Human cortical organoids (hCOs) were derived through the directed differentiation and morphological induction of human-induced pluripotent stem cells (hiPSCs). The integration of hCOs with microelectrode array (MEA) chips established hybrid bio-electronic agents. Within this framework, multiple hybrid agents interact in a closed-loop paradigm, each assigned distinct functional roles and tasks, collectively forming the MAIS architecture. The system is applied to neurological disorder research, wherein “doctor agents,” “patient agents,” and “control agents” are constructed via healthy and disease-model hCOs. Through closed-loop perturbation within the MAIS framework, targeted perturbative effects on pathological neuronal activity are achieved. **b)** Architecture of closed-loops. The hCOs are interfaced with MEA chips via a silicon substrate, facilitating bidirectional communication between the biological components and electronic systems. Neuronal signals generated by the hCOs are captured by the MEA chip and transmitted to an external computation module for processing. Pre-programmed instructions within the computation chip are then relayed back to the MEA chip to deliver targeted stimulation to the hCOs, completing a real-time feedback loop. **c)** Design of spatial reinforcement learning within MAIS. A human-interpretable reinforcement learning paradigm is implemented, combining card game analogies with neuronal electrical stimulation. In this framework, the hCOs functions as the agent, while the MEA chip and computation module constitute the environment. Electrical signals generated by the hCO during interactions represent “actions,” while stimuli dispatched from the MEA chip and computation module are defined as “states.” Differentiated stimulation signals (reward and punishment) from the MEA and computation chip are mapped to card ranks within the game paradigm. The reinforcement learning process facilitates adaptive perturbation of hCOs activity based on predefined reward-punishment criteria derived from the card game outcomes. **d)** Multi-agent modulation within MAIS. The MAIS framework extends the single-agent design to encompass multi-agent interactions. Neuronal activities of hCOs are represented and perturbed by reciprocal electrical stimulation within a shared closed-loop system. The “doctor agent” integrates neuronal signals from both the “patient agent” and the “control agent,” performing comparative analyses to determine the appropriate reward or punishment to administer. This process aims to recalibrate the “patient agent’s” neuronal activity to emulate the normative activity levels observed in the “control agent,” achieving dynamic perturbation within the multi-agent system.

## Result

### Characterization of Features of MAIS

The MAIS platform establishes a bidirectional interface between biological neuronal networks and synthetic electronics through its closed-loop circuit design, enabling real-time modulation of functional neuronal Activity. The biological module was constructed using our published cortical organoid (hCO) differentiation protocol^29^. The time-dependent variation of MAIS characterization starting from the induction of human pluripotent stem cells (hPSCs) into cerebral organoids (Day 1) was presented as a time-line in Fig. 2a. After a 50-day incubation, the co-staining of hCOs with SIM312 and MAP2 revealed extensive neurite outgrowth in the hCOs, implicating the establishing of complex neuronal networks, and shown in Fig. 2b. Sequential slicing of hCOs and mounting on HD-MEA chips was completed on Days 55 and 60, creating the construction of MAIS (Fig. 2c and Extended Data Fig. 1). Single-nucleus RNA sequencing (snRNA-seq) at Day 80 identified cellular heterogeneity, encompassing progenitor populations such as radial glial cells (RGCs) and intermediate progenitor cells (IPCs), as well as neuronal subtypes including newborn neurons (NNs), deep-layer neurons (DPNs), and upper-layer neurons (UPNs). Mature neurons, i.e., DPNs and UPNs, constituted 52.58% of the total cell population, which presented in Fig. 2d. Transcriptomic profiling verified the enrichment of cortical-specific genes, whilst immunofluorescent staining illustrated the self-organizing spatial distribution of cortical cell populations, showing consistency between the two results (Extended Data Fig. 2a-d). Longitudinal imaging from Day 30 to 80 further revealed progressive expansion of cortical-specific neurons during hCO maturation (Extended Data Fig. 2e-i). Gene ontology analysis highlighted remarkable upregulation of pathways associated with neuronal function in the neuronal population (Fig. 2e). The existence of synapse was validated at Day 120 through co-localized pre- and postsynaptic protein markers (Fig. 2f). In summary, these results demonstrate the establishment of functional neuronal networks within MAIS-integrated hCOs, providing evidences of neuronal connectivity for bidirectional bioelectronic interfacing.

**Figure 2.**
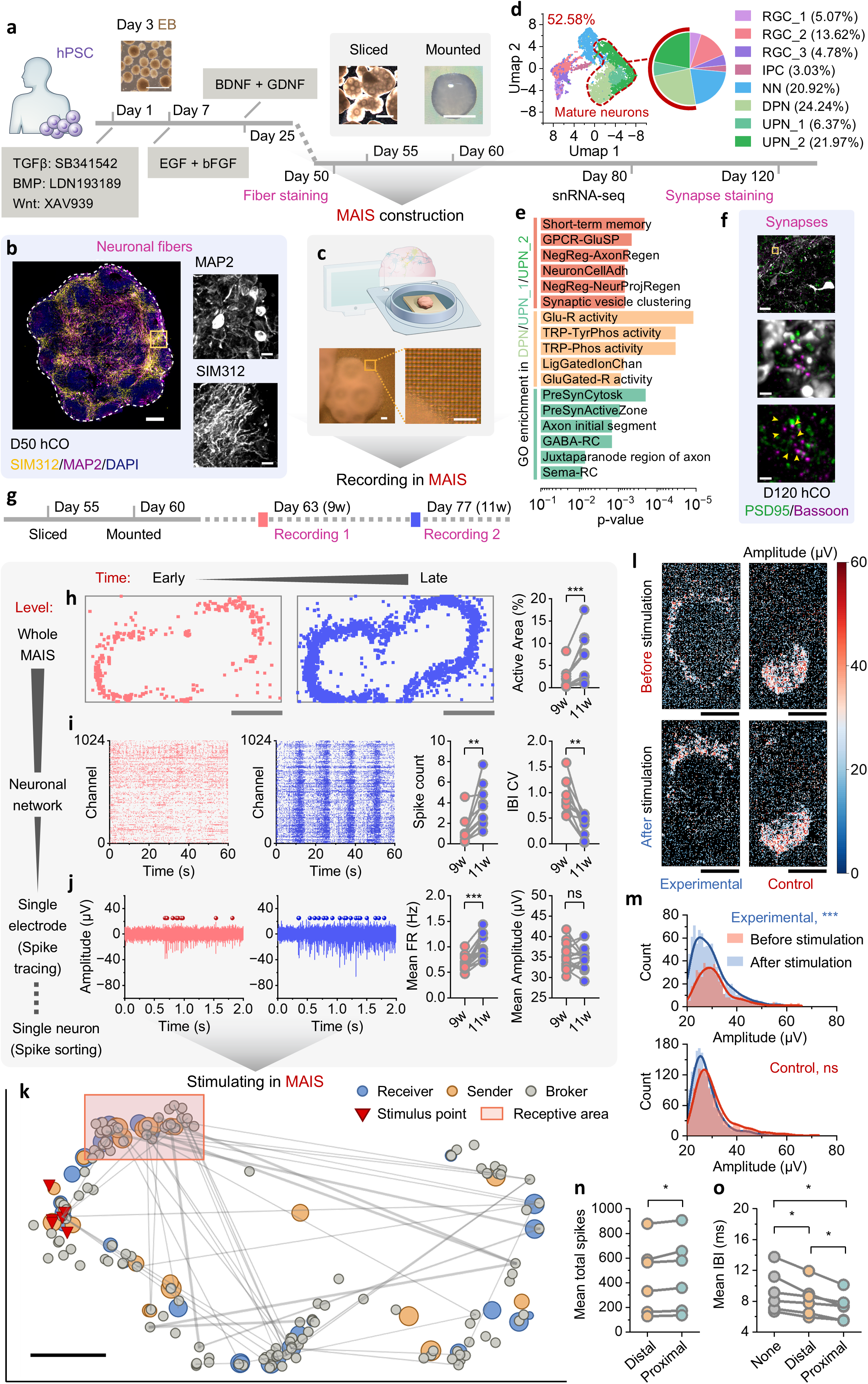
Characterization of Features of MAIS. **a**) Schematic of the differentiation process of brain organoids, along with representative bright-field images of midbrain organoids cultured in vitro. Scale bar, 500 μm. **b)** Immunostaining for SIM312 (axons) and MAP2 (dendrites) antibodies in Day 50 hCO within MAIS. The inset shows typical SIM312+ fibers and MAP2+ cells. Dashed line indicates the border of hCO. Scale bar, 100 μm. **c)** Schematic of the coupling between hCO and the MEA Maxone electrophysiology as MAIS, depicting localized coupling between the hCO and MEA chip as observed under bright-field microscopy. **d)** GO enrichment analysis of upregulated genes in the DPN and UPN cell populations from the snRNA-seq results of 80-day-old organoid cultures, highlighting the top 5 enriched GO terms. **e)** UMAP dimensionality reduction of snRNA-seq results from 80-day-old organoid cultures, with the pie chart on the left indicating the percentage of annotated cell populations. RGC: Radial glial cells; IPC: Intermediate progenitors; NN: Newborn neurons; DPN: Deep-layer neurons; UPN: Upper-layer neurons. **f)** Immunostaining for PSD95 (postsynaptic marker) and Bassoon (presynaptic marker) in Day 120 hCO within MAIS. Arrows indicate typical PSD95+ and Bassoon+ signals, suggesting the presence of potential synapses. Scale bar, 15 μm; 2 μm (inset). **g)** A timeline indicating the durations for slice culture and the placement of organoid cultures onto the MEA chip. Recording 1 represents signal collection for day 63 (9 weeks) and recording 2 represents signal collection for day 77 (11 weeks). **h)** Whole-array level active area mapping of sliced hCO within MAIS. Visualization of the active area regions in 9weeks (9w, red) and 11weeks (11w, blue) hCO samples. Based on whole-array HD-MEA activity scanning of hCO, the active area is defined as firing rate > 0.2 Hz and spike amplitude > 20 μV. The differences in active areas between paired samples of 9w and 11w hCO were quantified (n = 14), analyzed via the Wilcoxon matched-pairs signed-rank test, ***, p = 0.0001. **i)** Visualization of multi-electrode activity at the neural network level. Differences in spike per burst per electrode and coefficient of variation of inter-burst intervals (IBI CV) between paired samples of 9w and 11w were quantified (n = 10) using the Wilcoxon matched-pairs signed-rank test, with p = 0.002 for spike per burst per electrode and p = 0.002 for IBI CV. **j)** Spike tracing at single electrode level. hCOs within MAIS exhibited a range of network burst patterns, including activity resembling spindle bursts. The two panels show the aggregated multi-unit activity (0.5-3 kHz) recorded from the same electrode at 9w (red) and 11w (blue). The difference of the mean FR of paired samples of 9w and 11w by Wilcoxon matched-pairs signed-rank test (n = 14), ***, p = 0.0002. Difference of mean amplitude of paired samples of 9w and 11w by paired t-test (n = 14), ns, p = 0.7051. **k)** Visualization of the selection principle for transmitter and receiver signal regions via the Spike Timing Tiling Coefficient (STTC) method. Dimensionality reduction is performed on the electrical activity of hCO, followed by correlation analysis among neurons via the STTC algorithm, confirming the existence of signal transmission and reception correlations among different neurons. A subset of neurons with transmission capability (send) is designated as the sending area, while a subset with reception capability (receive) is designated as the receiving area. Selection diagram of transmitter and receiver signals in hCO based on STTC calculations. A correlation threshold of 0.22 was applied. **l)** Visualization of the impact of electrical stimulation on the amplitude of hCO within MAIS under the same differentiation conditions and cultivation duration. Experimental group: with electrical stimulation; control group: without electrical stimulation. **m)** Statistical analysis of the impact of electrical stimulation on the amplitude of hCO within MAIS under the same differentiation conditions and cultivation duration. The Kolmogorov-Smirnov (K-S) test reveals no significant difference in amplitude distribution for the control group (ns, p = 0.463), while the experimental group shows a significant difference (***, p = 0.0001). **n)** Experimental statistical results validating the spatial distance modulation effect of STTC stimulation-based spatial design in MAIS system on signal reception differences in the receptive area. Under stimulation conditions, the effects of proximal (closer to STTC connection) versus distal (farther from STTC connection) stimulation on mean total spikes in receptive area were quantified, showing significant differences between proximal and distal stimulation (Wilcoxon matched-pairs signed-rank test, n=6, p=0.0313). **o)** Experimental statistical results validating the spatial distance modulation effect on network-level electrical activity intensity in MAIS neuronal networks. Effects of proximal stimulation, distal stimulation, and none (control) on mean IBI in MAIS were quantified. Significant differences were observed between: proximal vs. distal stimulation (Wilcoxon matched-pairs signed-rank test, n=6, p=0.0313); distal stimulation vs. none (p=0.0313); and proximal stimulation vs. none (p=0.0313).

Grounded in the biological foundation of the MAIS-integrated functional neuronal networks, systematic electrophysiological characterization of network dynamics was further proceeded. Spatiotemporal activity analyses were conducted comparatively, focusing on the developmental phases at week 9 (Day 63, Recording 1) and week 11 (Day 77, Recording 2), and the experimental time-line was displayed in Fig. 2g. Regarding the whole-array resolution, full MEA scans indicated significant expansion of spontaneously active areas between weeks 9 and 11 (Fig. 2h, n = 14; ***P = 0.0001), suggesting that our MAIS achieves functionally stable interfaces within three weeks of coupling, accompanied by a gradual emergence of network maturation. For network-level resolution, we analyzed the synchronized activity from the top 1,020 active electrodes during a 300-s recording period, presenting 60-s data as an example in Fig. 2i. Quantitative comparisons of network synchrony metrics between week 9 and week 11 revealed two key maturation signatures of functional network: enhanced burst complexity through increased spikes per burst (n = 10; **P = 0.002) and stabilized network synchrony evidenced by the reduced inter-burst interval (IBI) variability (n = 10; **P = 0.002). The availability of multiple maturation indexes collectively indicates the progressive development of coordinated network bursting patterns. The temporal refinement of neuronal activity was further demonstrated by single-electrode resolution analysis in Fig. 2j (n = 14). Spike tracing showed significantly elevated in firing frequency at week 11 compared to week 9 (***P = 0.0001), whereas spike amplitudes remained stable (ns, P = 0.7051), such frequency-specific variations suggesting a promising synaptic maturation. The comprehensive electrophysiological profiling across multiple scales uncovers three fundamental principles governing the maturation of MAIS networks: the spatiotemporal expansion of spontaneous activity, the emergence of coordinated network-level bursting patterns, and the progressive refinement of single-unit firing properties. These findings show that MAIS facilitates the bidirectional bioelectronic integration where the progressive maturation of biological network translates into improved functional signal coordination at the bioelectronic interface.

After conducting a multiscale analysis of the spontaneous appearance of network dynamics during MAIS maturation, we further developed an interpretable stimulation protocol guided by intrinsic spatiotemporal electrophysiological correlations. Employing Kilosort2 for spike sorting, in conjunction with single-unit activity analysis (refer to Extended Data Fig. 3), resulted in the creation of detailed maps depicting pairwise spike correlations and the corresponding latency distributions^30^. The functional connectivity profiles unveiled the network architectures that emerge from the intrinsic neuronal coupling mechanisms. Spike time tiling coefficient (STTC) analysis identified the temporally coordinated patterns between recording units^31^, using a correlation threshold of 0.4 to visualize the dominant functional connections (Fig. 2k). Subsequent latency distribution analysis uncovered directional signaling architectures consistent with the fundamental axonal conduction principles^32,33^. Neuronal subpopulations that consistently initiate signals within a latency of less than 5 ms post-stimulus were operationally designated as Senders, while those showing largely delayed response latency, exceeding 15 ms, were classified as Receivers. This spatiotemporal classification adheres to established functional circuit mapping paradigms where latency gradients delineate the roles of efferent and afferent network^12,34^.

Building on this latency-based efferent-afferent functional mapping framework, a systematic linkage of endogenous spontaneous activity patterns in cerebral organoids with externally targeted neuromodulation, mediated through evidence-based circuits, can be realized by selecting spatial regions of high Sender density as stimulus points, and by defining Receiver-rich regions as receptive areas. Comparative analysis between electrically stimulated (Experimental) and non-stimulated (Control) groups under identical culture conditions have shown stimulus-specific network modulation. While the control groups maintained stable amplitude distributions pre-/post-intervention through Kolmogorov-Smirnov (KS) test (ns, P=0.463), the experimental groups displayed a notable increase in amplitude of post-stimulation (KS test, ***P=0.0001; Fig. 2l&m), indicating that the potentiation of MAIS amplitude dynamics was induced by stimuli, facilitated by circuit engagement guided by connectivity derived from STTC. To investigate spatial determinants of stimulation efficacy, we newly developed a 100-cycle alternating protocol comparing proximal versus distal stimulation relative to receptive areas. Stimulation distance-dependent quantitative analysis demonstrated the significantly greater response-evoked amplitudes in receptive areas following proximal stimulation compared to distal stimulation (KS test, *P=0.01). These functional connectivity-guided neuromodulation findings revealed two facts: STTC-derived connectivity mapping guides functional stimulation target selection through endogenous network architecture analysis, and spatial conditioning parameters are crucial in determining neuromodulatory efficacy. The described spatial modulation capacity provides essential operational principles for implementing closed-loop reinforcement learning in MAIS.

### Spatial reinforcement learning

The MAIS platform employed spatially organized bioelectronic reinforcement protocols within a closed-loop architecture in the “Card Games” paradigm as shown in Fig. 3a. The architectural components of the closed-loop system comprised three core modules: Stimulus encoding, involving a spatial patterning of 4 ms/50 mV through designated electrodes mapped to card values (Fig. 3a i-ii); Response decoding, which entails real-time neural computation based on the quantification of spike numbers within 1-s temporal windows (Fig. 3a iii); Adaptive feedback, featuring fixed-pattern rewards versus stochastic punishments contingent on decision accuracy (Fig. 3a iv). Electrode dynamics across task phases were quantified through firing rate analyses and spike count visualizations in Fig. 3b. Reward conditioning implemented the long-term potentiation (LTP)-like protocols^35–37^ using precisely timed (a 4-s interval), spatially coordinated (24-cluster activation) 50 Hz pulse trains (50 mV, 4 ms/phase) to elicit LTP-like plasticity through temporal coherence, synchronous spatial activation, and suprathreshold intensity. Conversely, punishment conditioning disrupted network synchrony through stochastic electrical stimulation (10-3000 ms intervals; 10-300 mV amplitude) characterized by temporal unpredictability, dynamic amplitude modulation, and asynchronous cluster activation. Notably, the fixed-intensity stimulus parameters in reward protocols potentially induced sustained depolarization that may facilitate spike-timing dependent plasticity (STDP) mechanisms^38,39^ through time-modulated neuronal interactions, whereas the variable stimulation parameters during punishment produced controlled synaptic noise patterns mimicking inherent network instability via stochastic desynchronization. This parametric dichotomy yielded opposing plasticity effects (Fig. 3c), i.e., reward stimuli decreased 90th percentile (P90) interspike intervals (Fig. 3d, reducing from 110.2 to 104.4 ms; Wilcoxon test, n=5, *p=0.031) potentially showing LTP-like properties, while punishments increased P90 intervals (Fig. 3d, raising from 103.4 to 106.9 ms; Wilcoxon test, n=5, *p=0.031) potentially indicative of long-term depression (LTD). The bidirectional modulation aligns with Hebbian principles—synchronous activation enhances connectivity, while asynchronous stimulation induces synaptic depression^40^.

**Figure 3.**
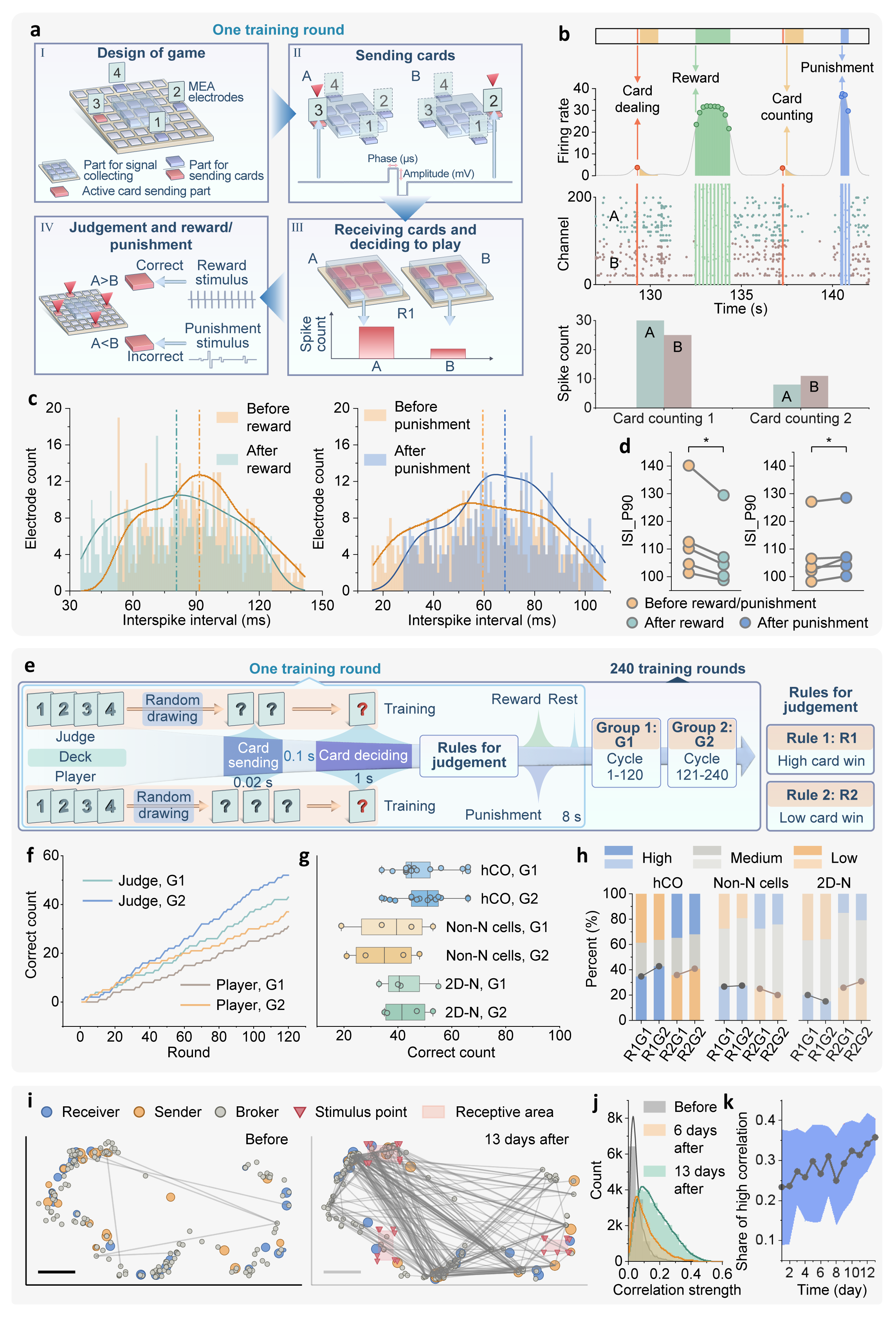
Spatial reinforcement learning. **a**) Schematic of MAIS closed-loop architecture in “Card Games” paradigm: i) stimulus encoding with 4 ms/50 mV spatial patterning mapped to card values. ii) electrode activation mapping. iii) response decoding via 1-s window spike quantification. iv) adaptive feedback of reward stimulus and punishment stimulus with fixed/stochastic conditioning. **b)** Electrode dynamics quantification showing firing rate distributions and spike count visualizations across task phases. **c)** Interspike interval (ISI) modulation under behavioral conditioning: i) reward protocols induced leftward ISI distribution shift ii) punishment protocols induced rightward shift. Data from representative hCO sample (n=1) displayed through histogram bins with superimposed KDE curves. Dashed vertical lines mark distribution means. **d)** Plasticity modulation showing reward-induced P90 interval reduction (110.2→104.4 ms, *p=0.031) vs punishment-induced increase (103.4→106.9 ms, *p=0.031). **e)** Role-specific training framework with protocol implementation: i) single-round procedure with judges receiving 2 cards from a 4-card deck and players receiving a 3-card sequence. The duration of the card dealing lasts 0.02s. ii) decision-making protocol including spike count quantification initiated 0.1s post-stimulus in receptive regions, 1-s temporal window analysis for card value decoding and reward/punishment conditioning based on rule compliance. iii) performance progression in roles training: G1 (rounds 1-120) vs. G2 (rounds 121-240) showing optimization trends. iv) rule architecture including R1 with high card win as the value of the card higher than the opponent’s value and R2 with low card win as the value of the card lower than the opponent’s value. **f)** The cumulative line graph compares the cumulative changes in the correct times of the card game between the first 120 rounds (G1) and the last 120 rounds (G2) of the agent’s training. **g)** Protocol-driven optimization comparison: hCO-MAIS (+18.7% Δ), 2D cultures (+4.1%), and non-neuronal controls (−8.3%). Wilcoxon tests show non-significant trends (p=0.212-0.743, n=3). **h)** Rule adaptability through R1→R2 inversion: hCO-MAIS maintains directional optimization (+22.4% R1 vs +19.7% R2) vs stochastic responses in controls. **i)** Functional connectivity progression: STTC >0.4 connections increase from 0.01% to 2.31% (***p=0.0001, n=8). Spatial mapping shows stimulus-receptive area specialization. **j)** Network synchronization shift: correlation strength distributions pre- vs. post-training (13-day) comparison (KS test ***p=0.0001, n=8) **k)** Line plots (n=8) of the change in the proportion of high correlation (STTC >0.4 connections) with increasing number of days in the study.

The role-specific training framework, which defines two distinct neuronal network roles—Judges and Players—implements differentiated protocol loads based on task hierarchy illustrated in Fig. 3e. Judges were presented with two-card combinations per trial, mirroring their binary discrimination function, whereas Players dealt with sequences of three cards, which enabled the analysis of sequential pattern. Comparative analysis between the initial phase, i.e., G1 for rounds 1-120, and advanced phase, i.e., G2 for rounds 121-240, of Judge training revealed progressive enhancement in performance (Fig. 3f). The differences in protocol-driven optimization were validated by diverse cellular experiments and are plotted in Fig. 3g. hCO-MAIS exhibited non-significant but directionally consistent improvement from G1 against G2 in performance (defined as Δ; +18.7%, p=0.212; Wilcoxon test, n=3), suggesting tentative protocol-driven optimization. In contrast, non-neuronal controls (Δ ∼ -8.3%, p=0.498, n=3) and 2D neuronal cultures (Δ ∼ +4.1%, p=0.743, n=3) showed no directional trends. The upward performance trends consistently observed in all hCO-MAIS replicates (n=3) warrants further investigation beyond the current statistical parameters. Notably, the observed architectural specificity—with hCOs showing superior task-oriented optimization compared to monolayer cultures— suggests that three-dimensional cytoarchitecture may enhance adaptive plasticity in biohybrid systems.

Rule adaptability was examined through protocol inversion, i.e., shifting from rule R1 (the high-card win protocol) to rule R2 (the low-card win protocol). hCO-MAIS exhibited consistent directional optimization across both selection regimes, increasing Δ by +22.4% under high base selection (R1, p=0.114) and +19.7% under low base selection (R2, p=0.158), in spite of not reaching statistical significance for either difference (Fig. 3h). The observation of insignificant direction maintenance effects possibly originates from protocol-dependent synaptic inertia requiring extended training phases for rule reconsolidation, cross-regulatory interference between competing reinforcement protocols, and non-linear scaling of decision complexity in inverted rule space. Both Non-N and 2D-N groups exhibited stochastic response characteristics devoid of systematic directional bias, showing high inter-regime variability (R1 vs. R2: variance exceeding 25% for both groups), congruent with non-directional fluctuation patterns. These findings suggest that MAIS preserves adaptive potential within predefined artificial rule architectures, though achieving robust decision transformations in constrained experimental frameworks (e.g., our card-matching paradigm) may require optimized training durations and protocol decoupling strategies. The observed rule-sensitive adaptation trajectories provide an example for developing context-aware bioelectronic hybrid decision systems.

Expanding on the observed performance optimizations, the temporal and spatial reorganization of network connections during MAIS training was characterized and shown in Fig. 3i-k. Functional connectivity analysis showed a gradual increase in strongly synchronized connectivity across 8 specimens, with the proportion of STTC >0.4 correlations rising from 0.01% (6/59,340) pre-training to 2.31% (1,430/61,776) after 13 days (Wilcoxon test, ***P=0.0001; Fig. 3i). This significant enhancement coincided with two complementary phenomena, including a training-dependent shift in network synchronization patterns manifested by rightward displacement of correlation strength distributions after 13-day training (KS test, ***p=0.0001, n=8; Fig. 3j), and progressive expansion of functional connectivity evidenced by linear increases in the proportion of high-correlation connections (STTC >0.4) across experimental days as shown in Fig. 3k. Spatial mapping, performed in Fig. 3i, revealed that neuronal nodes operating in synchrony (Receivers, Senders, and Brokers) predominantly establish connections between specific stimulus points and their corresponding receptive areas, reflecting the spatial modulation logic inherent in our closed-loop paradigm. These results demonstrate two significant insights: spatially constrained reinforcement may potentially strengthen the connectivity of stimulus-receptive regions; the proliferation of strongly synchronized connections (from 6 to 1,430 connections with STTC >0.4) correlates temporally with achieving decision transformations in constrained experimental frameworks, although this correlation requires further causal validation. By integrating the temporal hierarchy of plasticity — spanning millisecond-scale LTP-like dynamics, daily connection increments, and multi-week network remodeling — these findings collectively establish MAIS as a configurable platform capable of characterizing the decision-making behaviors of bioelectronic interfacing in constrained frameworks and probing neuroplasticity dynamics across spatiotemporal scales through its single-agent implementation, while providing foundational architecture for scaling to multi-agent systems.

### Neuronal dynamics across multi-agent interactive system

Expanding from the reliable single-agent system in Fig. 3, our MAIS platform progresses through three developmental stages: initially establishing biological decision module in Solo Mode, subsequently constructing competitive frameworks with rule-based adjudication in Dual Mode, and ultimately achieving autonomous biological competition-adjudication cycles in Triple Mode. This architectural evolution systematically advances from elementary neural decision-making to full biological resolution of multi-agent hierarchies (Fig. 4a). Solo Mode (1P) forms the fundamental module where a single hCO player demonstrates deterministic card selection through maximal spiking activity. This closed-loop unit establishes the elementary biological decision mechanism through stimulus-response mapping, serving as the building block for multi-agent interactions. Dual Mode (2P) constructs the competitive framework by introducing biological competition between two hCO players (P1/P2). The system implements rule R1-based computer adjudication to resolve decision hierarchies - where card magnitudes determine agent dominance levels. This mode establishes the critical “compete-adjudicate” paradigm that bridges single-agent decisions with multi-agent interactions. Triple Mode (3P) achieves full biological adjudication by incorporating a third hCO judge that replaces computerized arbitration. The competing players’ card selections (decision outputs) are converted into spatiotemporal stimulus patterns for the judge organoid, whose differential spiking activity directly determines outcome allocation: Higher activity at Position B (representing P2’s input) triggers reward-penalty assignments, establishing endogenous biological hierarchies through agents competition. This progression demonstrates our platform’s unique capability to simulate multi-agent decision-making systems where both competitive actions and hierarchical resolutions emerge from biological neural processes.

**Figure 4.**
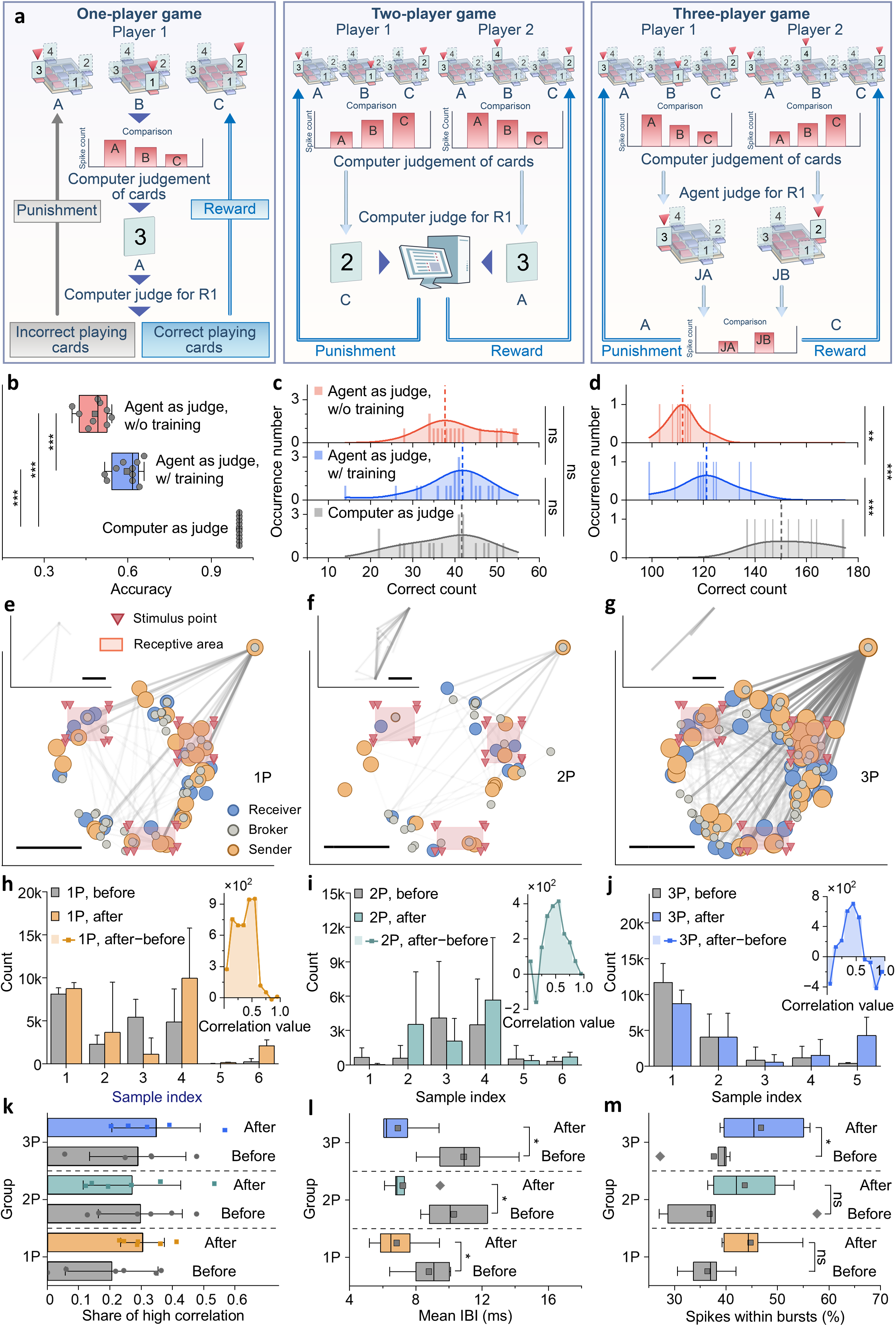
Neuronal dynamics across multi-agent interactive system. **a**) The card game system evolves through three operational tiers: In One-player-game (Solo Mode), a single hCO interfaces with a microelectrode array (MEA) as player/judge, where maximal spiking activity at designated positions (A/B/C) determines card selection (e.g., Position A corresponds to Card 3), with Rule R1 validation triggering reward signals to active electrodes. Expanding to Two-player-game (Dual Mode), two hCOs players (P1 and P2) competitively generate position-specific spiking patterns (e.g., P1’s Position C corresponds to Card 2 vs. P2’s Position A corresponds to Card 3), where computational arbitration under R1 delivers differential conditioning (reward/punishment) based on card hierarchy. Further advancing to Three-player-game (Triple Mode), a third hCO replaces computerized judgment: P1/P2’s card selections are encoded as patterned stimulation to the judge organoid, whose competitive spiking at input-specific positions (A/B) directly governs outcome allocation - higher activity at Position B (representing P2’s input) triggers P2’s reward and P1’s punishment, establishing biological adjudication of decision hierarchies through emergent neuronal competition. In the experimental setup, Solo Mode serves as the basis for Triple Mode, with a 2-hour interval between different modes. **b)** The judge efficiency comparison illustrates the judgment accuracy of three modes in game decision-making: the Dual Mode with computer as judge, Triple Mode with agents as judge trained in Solo Mode (n=6), Triple Mode with agents as judge untrained in Solo Mode (n=7). K-S tests revealed statistically significant pairwise differences: computer as judge vs. trained agent as judge(**P<0.001), computer as judge vs. untrained agent as judge (**P <0.001), and trained agent as judge vs. untrained agent as judge (*P <0.05). **d)** Different judge participation scenarios were compared by overall correct count distributions. Computer as judge showed significant differences in distribution of correct counts compared to both untrained agent (**P <0.001) and trained agent (**P <0.001) via the K-S test. A significant difference (*P <0.05) was also observed between untrained agent and trained agent. **e-g)** Post-game neuronal dimensionality reduction analyses (threshold=0.7) show enhanced functional connectivity across modes (Solo/Dual/Triple). All modes exhibited increased highly correlated connections (>0.7) after stimulation. The topological reorganization demonstrates experience-dependent plasticity in hCO networks, with Triple Mode revealing complex competition patterns through position-specific connectivity emergence. **h)** Analysis of pre-/post-Solo Mode correlation distribution (n=6) reveals connectivity enhancement: five samples showed increased counts across all correlation bins (0-1). Pooled data revealed maximum enhancement in mid-range correlations (0.1-0.6, Δ>695 connections). **i)** Dual Mode induced moderate connectivity growth (n=6): three samples showed universal enhancement, with cumulative increases strongest in the 0.3-0.6 range (Δ>328). **j)** Triple Mode induced connectivity restructuring (n=5): Bidirectional changes occurred with 0.6-1 correlations decreasing (0.8-1, Δ>199) while 0.3-0.6 connections increased (Δ>522). **k)** Resting state high correlation (STTC>0.5) ratio comparisons revealed mode dependent plasticity: Solo mode showed maximal enhancement, Triple mode showed moderate enhancement, and Dual mode showed slight reduction. **l)** Inter-burst interval (IBI) analysis showed universal modulation across modes (Solo p=0.0312, n=6; Dual p=0.0312, n=7; Triple p=0.0312, n=6). All modes significantly shortened post-game IBIs. **m)** Burst pattern analysis revealed differential modulation: Solo Mode significantly increased the proportion of spikes within bursts (*p=0.0312), while Dual (ns, p=0.07) and Triple Modes (ns, p=0.0625) showed non-significant trends.

The comparative efficiency analysis of adjudication systems revealed differences between computational and biological decision-making. Although the computer judge showed statistically superior accuracy compared to both trained (Fig. 4b; K-S test, *P<0.001) and untrained hCO judges (K-S test, **P<0.001), the difference in performance between trained and untrained biological judges (K-S test,*P<0.05) suggests a capacity for adaptive learning within MAIS’s biological modules. This finding is further substantiated by the distinct distributions of correct decisions across different groups of judges, as revealed through progressive analytical perspectives (Fig. 4c and d). In the player performance analysis (Fig. 4c), comparisons of player performance between computer judge group versus both trained agent judge group (K-S test, ns, P>0.05) and untrained agent judge group (K-S test, ns, P>0.05) showed no statistically significant differences in players’ correct decision distributions. Similarly, no distinction of player performance emerged between trained and untrained biological judges groups (K-S test, ns, P>0.05). In contrast, the integrated player-judge system analysis (Fig. 4d) revealed significant disparities: the integrated performance of player-judges in computer judgment group differed markedly from both agent-judge conditions groups (K-S test, **P<0.001), while trained biological judges outperformed untrained counterparts (K-S test, *P<0.05). This tripartite analytical framework systematically dissects the multi-agent interaction efficiency within MAIS through pure judge performance (Fig. 4b), isolated player outcomes (Fig. 4c), and coupled player-judge dynamics (Fig. 4d), progressively unveiling how the agent components of MAIS shape decision-making efficacy in agent competition paradigms.

Comparative analysis of post-task neuronal connectivity revealed distinct patterns of network reorganization patterns across game modes (Fig. 4e-g). Dimensionality reduction analyses (threshold=0.7) demonstrated universal post-stimulation enhancement of highly correlated connections (>0.7) across all modes. Solo Mode induced maximal connectivity enhancement (n=6), with five experimental replicates showing increased connections across all correlation bins (0-1), peaking at intermediate correlations (0.1-0.6, Δ>695; Fig. 4h). Dual Mode showed moderate enhancement (n=6), with increases observed in three samples and the strongest cumulative increases in 0.3-0.6 correlations (Δ>328; Fig. 4i). Triple Mode (n=5) revealed bidirectional restructuring - decreased high correlations (0.6-1, Δ>199) contrasted with increased 0.3-0.6 connections (Δ>522; Fig. 4j). Resting state analysis (Fig. 4k) confirmed mode-dependent plasticity: Solo Mode showed a maximal increase. Triple Mode exhibited a moderate increase. And Dual Mode demonstrated a slight decrease. The observed connectivity divergences may reflect different rule implementation: deterministic reinforcement of Solo Mode (single-player, error-free adjudication) allows for consistent network strengthening, while Dual Mode’s intermittent stimulation (potential draw outcomes reducing conflict resolution demands) may attenuate consolidation efficiency, whereas Triple Mode’s probabilistic feedback (hCO-mediated stochastic rule application) requires dynamic filtering - strengthening task-relevant mid-range correlations (0.3-0.6) while pruning non-adaptive high correlation connections (>0.6). This hierarchy aligns with the certainty gradients of rules across experimental paradigms. The quantitative changes align with subsequent electrophysiological features -particularly temporal reorganization of inter-burst intervals (Fig. 4l) and intra-burst spiking dynamics (Fig. 4m), forming a multi-dimensional evidence chain that deciphers spatiotemporal coding characteristics in the decision-making processes of MAIS.

Complementary electrophysiological characterization revealed mode-specific temporal patterns. Marked differences in the inter-burst interval (IBI) are evident across all modes (Fig. 4l), as opposed to the changes of spikes within bursts that are exclusively observed in Solo Mode (Fig. 4m), suggesting distinct multi-scale encoding hierarchies between individual and interactive decision-making scenarios. This multi-scale quantification framework combines two critical biological metrics: cellular-level electrophysiological signatures revealing temporal bursting patterns, and network-level interaction dynamics mapping neuronal correlations. Together, these metrics establish the capability of MAIS to decode emergent biological computations across hierarchical levels: from single-agent neural encoding to multi-agent coordination.

## Discussion

Our work establishes MAIS a biologically interpretable in vitro neural modulation framework by leveraging the inherent self-organizing principles and synaptic plasticity of neuronal networks in cortical organoids. Traditional in vitro systems achieve specific encoding and decoding strategies by superimposing additional artificial rules^22–24^, where artificially defined spatiotemporal regulations force biological networks to operate under static predefined templates over extended periods, thereby compromising the inherent self-organizing encoding and decoding features carried by dynamic neuronal networks. MAIS fundamentally rethinks this paradigm through plasticity-aware stimulation -encoding intervention signals not as preset static patterns, but as spatiotemporal sequences that integrate endogenous synaptic remodeling processes. This neuronal network modulation strategy emphasizing the intrinsic complexity of neuronal networks in MAIS enables the transition from single-agent programming to multi-agent coordination while preserving biological interpretability, demonstrating that neuronal network regulation can be systematically addressed through adaptive spatiotemporal encoding rather than mechanical control. The adaptive spatiotemporal encoding first proposed in MAIS grants human-derived neuronal networks greater autonomy during long-term closed-loop modulation within in vitro models, while the preservation of intrinsic biological complexity within the MAIS framework enables deeper investigation into the encoding strategies of human-derived neuronal networks.

Current neural interface paradigms face fundamental limitations in scaling biological complexity. Firstly, predefined spatiotemporal architectures struggle to accommodate the dynamic topological evolution inherent in self-organizing biological networks, particularly when confronting the intrinsic variability of individual neural specimens^41,42^. Furthermore, conventional systems falter during multi-agent integration, as their artificial homogeneity fails to address biological heterogeneity while maintaining regulatory stability across scaled implementations^3,43^. The transition from single-agent to multi-agent architectures reveals a critical scaling principle: computer-guided single-agent systems governed by fixed spatiotemporal rules operate like shaping playdough with cookie-cutter molds - enforcing neuronal activity into rigid templates that demand exhaustive redesign for new component integration and fundamentally resist adaptation to evolving functional networks. Unlike conventional systems requiring exhaustive parameter tuning, MAIS exploits neuronal networks’ inherent capacity to assimilate stimulation into their functional logic - enabling coordinated multi-agent interactions through shared plasticity rules rather than centralized control. By preserving the autonomy of individual agents while establishing plasticity-guided communication protocols, MAIS transcends the limitations of both artificial rule-designed neuronal networks and traditional biological models - creating a sandbox interface where biological intelligence undergoes closed-loop evolution through self-driven plasticity. This paradigm not only advances our capacity to study complex neuronal dynamics in vitro but provides new perspectives for designing biohybrid systems, offering both foundational principles and testing scenarios for next-generation neural interfaces that genuinely respect biological complexity through engineering implementations like MAIS.

## Methods

### Stem cell line maintenance

hES cell lines H9 (WA09) was obtained from WiCell. The induced pluripotent stem (iPS) cell lines were generated using the Sendai virus carrying the Yamanaka reprogramming factors OCT3/OCT4, SOX2, c-MYC and KLF4. All cells were maintained in a 5% CO2 incubator at 37 °C and routinely tested for mycoplasma. All the cell lines were cultured under feeder-free conditions, seeded onto six-well plates coated with Matrigel (Corning) or Vitronectin (STEMCELL Technologies) and maintained in Essential 8 medium (Gibco). Cells were fed daily and passaged every 5-6 days using 0.5 mM EDTA solution treatment.

### Generation of human cortical organoids

Human hPSCs colonies on Vitronectin (5-6 days after passaging) were detached with dispase for 5 min at 37 °C. The colonies were suspended with Essential 8 medium for 24h to form organoids. Twenty-four hours following spheroids formation (day 1), medium was replaced by Essential 6 (Gibco) supplemented with 10µM SB431542 (MCE) and 100nM LDN193189 (MCE). In addition the Wnt pathway inhibitor 2.5 μM XAV-939 (MCE) was added. Medium were exchanged daily. On the seventh day in suspension, organoids were transferred to neural medium-A containing Neurobasal-A(Gibco), 1× N2 supplement(Gibco), 1× B27–vitamin A(Gibco), 1× GlutaMAX(Gibco), 1× MEM-NEAA(Gibco) and 1× antibiotic–antimycotic(Gibco). Neural medium was supplemented with 20 ng/ml EGF (R&D Systems) and 20 ng/ml FGF2 (R&D Systems) from day 6 to 20. Medium were changed every other day. To obtain purified neural patterning, spheroids were attached onto six-well plates from day 7 to day 13 to picking high-quality neural rosettes. On day 20, the organoids were cultured on an orbital shaker at a rotating speed of 50 rpm. On day 25, the medium was replaced with neural medium+A containing Neurobasal(Gibco), 1× N2 supplement, 1× B27(Gibco), 1× GlutaMAX, 1× MEM-NEAA and 1× antibiotic–antimycotic. From day 25 to day 39, neural medium was supplemented with 20 ng/ml BDNF (Peprotech) and 20 ng/ml GDNF (Peprotech), with medium change every other day. From day 40, only neural medium without growth factors was used for medium changes every 3 days.

### Cultured of Fibroblasts

Human fibroblasts were cultured in DMEM(Gibco) supplemented with 10% fetal bovine serum (Gibco)under standard conditions. Cells were used as a non-neural control and plated onto MEA as described below with the exception that testing began 48 h after plating as this cell type does not mature into electrically active cells.

### Dissociation of brain organoids

6 hCOs per hPS cell line were transferred to wells in six-well plates and incubated for 45–60 min at 37 °C with 3 ml of TrypLE (Gibco). Following incubation, samples were collected in a 15 ml Falcon tube with inhibitor solution and centrifuged at 300g for 2 min. The organoids were gently separated into single cells using p1000 pipette tip. Cell pellets were resuspended in culture medium consisting of Neurobasal supplemented with B-27 and 10 μM Y-27632. Undissociated tissue was removed by passing the cell suspension through a 40 μm cell strainer. Finally, dissociated cells were used for scRNA-seq or neuronal cell plating on HD-MEA.

### Sample preparation and immunofluorescent staining

Organoids were fixed in 4% paraformaldehyde for 2 hours and dehydrated with 30% sucrose solutions overnight. Organoids were serially sectioned at a thickness of 25 μm by frozen section (CM1950, Leica). All the samples were washed three times with PBS, then incubated in a blocking buffer (10% donkey serum and 0.3% Triton X-100 in PBS) for 60 min at room temperature before being incubated in the following primary antibodies overnight at 4°C. Fluorescently conjugated secondary antibodies were used to reveal the binding of primary antibodies (Jackson) and nuclei were stained with DAPI (sigma). Finally, the sections were washed and mounted to glass slides with Fluoromount-G. Images were captured on Olympus SpinSR confocal microscope.

Primary antibodies used were as follows: anti-SOX2 (mouse, R&D, MAB2018,1:1,000); anti-CTIP2 (rat, Abcam, ab18465, 1:1,000); anti-TBR1 (rabbit, Abcam, ab31940, 1:1,000); anti-SATB2 (goat, Santa Cruz, SC81376,1:500); anti-Bassoon (mouse, Abcam, AB82958,1:1,000); anti-PSD95 (rabbit, Abcam, ab18258); anti-MAP2 (rabbit, Santa Cruz, sc20172, 1:3,000); anti-SIM312 (mouse, Biolegend, 837904, 1:1,000); anti-PAX6 (rabbit, Biolegend, 901301, 1:1,000); anti-TBR2 (sheep, R&D, AF6166, 1:500).

Secondary antibodies used were as follows: Alexa Fluor 488 Donkey anti-mouse IgG (Invitrogen, Molecular Probe, A21202, 1:1,000), Alexa Fluor 594 Donkey anti-mouse IgG (Invitrogen, Molecular Probe, A21203, 1:1,000), Alexa Fluor 594 Donkey anti-rabbit IgG (Invitrogen, Molecular Probe, A21207, 1:1,000), Alexa Fluor 488 Donkey anti-rabbit IgG (Invitrogen, Molecular Probe, A21206, 1:1,000), Alexa Fluor 594 Donkey anti-goat IgG (Invitrogen, Molecular Probe, A11058, 1:1,000), Alexa Fluor 488 Donkey anti-rat IgG (Invitrogen, Molecular Probe, A21208, 1:1,000), Cy5 AffiniPure Donkey Anti-Goat IgG (H+L) (Jackson, 705-175-147,1:300), Cy5 AffiniPure Donkey Anti-mouse IgG (H+L) (Jackson, 715-175-150,1:300), Cy5 AffiniPure Donkey Anti-rabbit IgG (H+L) (Jackson, 711-175-152,1:300).

### Cellular quantification

To quantify the cellular population of NEUN, SOX2, TBR1, SATB2, CTIP2 expressing cells among total (DAPI-labeled) cells, stitching images under 60× objective lens from whole organoids were analyzed. Cells were automatically counted with IMARIS software.

### High-density microelectrode arrays

Human cortical organoids (hCOs) were recorded on single -well planar high-density microelectrode arrays (HD-MEAs) provided by MaxWell Biosystems (MaxWell Biosystems). The single-well complementary metal-oxide-semiconductor (CMOS)-based HD-MEA (“MaxOne”) comprises 26,400 platinum microelectrodes (electrode size: 9.3 × 5.3μm^2^) at a 17.5 μm pitch (centre to centre) within a total sensing area of 3.85 × 2.10 mm. The MaxOne system allows for simultaneous recordings from up to 1024 readout electrodes or channels at a sampling rate of 20 kHz. the electrodes can be flexibly selected and reconfigured according to experimental needs. For all setups, the acquisition filter band cut-off was approximately 300 Hz.

### Slicing and plating of hCOs on HD-MEAs

Cross-sectional slices were obtained from 7week-cultured hCOs. Whole organoids were embedded in 3% low-gelling-temperature agarose at ≈40 °C. Organoids embedded in the agarose gel were then sectioned into 360 μm thick slices with a vibratome (WPI). For recovery, slices were housed in transferred to ultralow-attachment plastic dishes, maintained in a 5% CO2 incubator at 37 °C. A recovery procedure of greater duration was designed, with a duration of one week, and a manual QC procedure was also added to the remove low-quality slices. The recovering medium containing Neurobasal, 1× N2 supplement, 1× B27, 1× GlutaMAX, 1× MEM-NEAA and 1× antibiotic–antimycotic, supplemented with 20 ng/ml BDNF (Peprotech), 20 ng/ml GDNF (Peprotech) and NT-3 (Peprotech).

To improve tissue adhesion, arrays were treated with 0.07% (v/v) poly(ethylenimine) (Sigma-Aldrich) in borate buffer for primary coating; and 0.04 mg/mL Laminin for secondary coating. Organoid slice was transferred to the HD-MEA well with a pasteur pipette and gently positioned over the recording electrode surface of the HD-MEA while visualizing with the stereoscopic microscope. After positioning the tissue, we used a small amount of Matrigel and several drops of recovering medium to enhance adhesion. Following the incubation, we carefully added more recovering medium. Half of the medium was changed every 2–3 days. During the first week, recovering medium was progressively replaced with recording medium. Recording medium prepared as follows: BrainPhys neuronal medium (StemCell Technologies), 2% (v/v) NeuroCultTM SM1 Neuronal Supplement (StemCell Technologies), 1% (v/v) N2 Supplement-A (StemCell Technologies), 1× GlutaMAX and 1× antibiotic–antimycotic, supplemented with 1 mM Dibutyryl-cAMP (Sigma), 20 ng/ml BDNF (Peprotech), 20 ng/ml GDNF (Peprotech) and NT-3 (Peprotech).

### Scanning electron microscopy

At designated endpoints, media was aspirated from the MEA wells and cells were fixed with fixative contain glutaraldehyde and paraformaldehyde for 12 h. They were then washed three times before being post-fixed with 1% OsO4 for 1 h. OsO4 was removed and the fixed cells were washed with three times in milliQ water and dehydrated via an ethanol gradient exchange (30%, 50%, 70%, 90%, 100%, 100% v/v) for 15 min each. After dehydration, the cells were dried and then allowed to evaporate. HD-MEA chips were manually cut to the appropriate size and sputter coated with 30 nm layer of gold. Coated MEA chips were then imaged using Hitachi SU8010 Ultra-high Resolution Scanning Electron Microscope.

### MK801 Synaptic inhibition experiment

We conducted stimulation experiments on organoid cultures at four conditions: baseline, +MK801 1h, washout 24h, and washout 48h. The parameters for the stimulation were set as follows: stimulation pulse phase at 100mV-1000mV with a 100mV step size, number of bursts = 10, and interpulse interval = 10ms. Statistical analysis was performed to evaluate the percentage of active electrodes, total number of spikes, and evoked peak for each of the four stages. The analysis was conducted with the following filtering parameters: firing rate threshold [Hz] < 0.2, amplitude threshold [μV] < 20, analysis window duration [s] = 0.5, and bin size [ms] = 50.

### Experimental setup and training of agents in a card game system

We developed a card game system using the MAIS platform to train agents to act as players and judges in the game, with three different roles: one judge and two players. Each agent corresponds to a different role. The card deck consists of eight unique cards (2 red1, 2 red2, 2 black1 and 2 black2). In each round of the game, the players randomly receive three cards from the deck and choose one as their play card. The judge, on the other hand, is responsible for judging whether the players’ chosen play cards comply with the rules of the game.

For example, in a game where the biggest card wins, the judge compares the two players’ play cards and determines which card is the biggest. The player who selects the largest card receives a reward stimulus, while the losing player receives a punishment stimulus. To increase training efficiency, the deck in training mode is simplified to only four unique cards (1 Red1, 1 Red2, 1 Black1, and 1 Black2). In this mode, the judge randomly receives two cards and conducts judgment training, while the players each receive three cards for judgment training. Each agent is trained independently.

During training, we define specific regions for card stimulation and response in each agent based on the distribution of active electrodes. As shown in Figure 3b, in the judge training scenario, two cards are assigned, resulting in two distinct response regions (corresponding to the A and B sections of the Sending Cards block in Figure 3b). For the player role, three cards are assigned, resulting in three different response areas. Each card response area is divided into a card stimulation area and a card stimulation information response area. The card stimulation regions are placed around their corresponding response regions.

Given the four different card face values, each card stimulus information response region covers four stimulus regions. We assign spatial locations to these card stimulation regions to differentiate the card values: the top portion is designated red2, the bottom portion is designated black1, the left portion is designated red1, and the right portion is designated black2, with the order of values being red2 > red1 > black2 > black1. During each training session, only one stimulation electrode is activated within each card response area, corresponding to a single card presented.

After the card stimulation is delivered, the total electrical signal within each card response area is recorded for 100 milliseconds, beginning 0.1 second after stimulation. The electrical signal response is then compared across regions, and the region with the highest signal is assigned as the correct playing card. If the selected card is correct, reward stimuli are delivered to all card stimulation regions. Conversely, if the selected card is incorrect, punishment stimuli are delivered to all card stimulation regions.

### Quantification and statistical analysis

Statistical analysis was carried out with Graphpad Prism software (9.5.1). Experimental data were presented as mean ± SEM. The normality of data distribution was evaluated with the Shapiro-Wilk test, and the homogeneity of variance was assessed by Levene’s test. When normality and homogeneity of variance tests were met, data were processed as follows: two-tailed Student’s t tests or ANOVA for two or multiple comparisons with one factor. When normality or homogeneity of variance failed, we used we used the Mann-Whitney U test for the comparison of two independent samples, Wilcoxon’s test for the comparison of two related samples. All details on sample size, the number of replicates, statistical tests and P values for each experiment are provided in the relevant figure legend. No statistical methods were used to pre-determine sample sizes owing to the exploratory nature of the experiments.

### Single-Nucleus Extraction Protocol for 10x Genomics Single-Cell Sequencing

The single-cell nuclear extraction was performed following the manufacturer’s instructions. We utilized the Nucleus Isolation Kit (SHBIO, Cat. No. 52009-10) for nuclear isolation. Briefly, tissue samples were first minced into small pieces (∼1 mm³) and transferred to a 1.5 mL tube containing ice-cold lysis buffer (10 mM Tris-HCl, 10 mM NaCl, 3 mM MgCl₂, 0.1% IGEPAL CA-630, and 1% RNase inhibitor, pH 7.4). Samples were gently homogenized using a Dounce tissue grinder (pestle A) with 10– 15 strokes on ice. The homogenate was incubated on ice for 5 minutes to ensure complete lysis of cell membranes while preserving nuclear integrity. The suspension was filtered through a 40 μm cell strainer into a fresh, pre-chilled 1.5 mL tube to remove debris. The filtered suspension was centrifuged at 500 × g for 5 minutes at 4°C to pellet the nuclei. The supernatant was carefully removed, and the nuclei pellet was resuspended in PBS supplemented with 1% bovine serum albumin (BSA) and 1% RNase inhibitor. Nuclei were stained with 0.1% Trypan Blue to assess viability and purity using a hemocytometer or automated counter. Approximately 20,000 nuclei were loaded onto a Single Cell G Chip (v3.1 chemistry, PN-1000120). The 10x libraries were constructed using the Chromium Single Cell Gene Expression Platform and Chromium Single Cell 3’ Reagent Kits (v3.1 chemistry, PN-1000121). In brief, single-nucleus suspensions were loaded into the channels of the Chromium Controller (10x Genomics, Pleasanton, CA) to generate single-cell GEMs (gel beads in emulsion). Library preparation was performed using the Chromium Single Cell 3’ Gel Bead and Library Kit v3.1 (PN-1000123, 1000157, 1000129; 10x Genomics). Libraries were sequenced on the Illumina Xplus platform.

## Acknowledgements

This study is supported by the National Natural Science Foundation of China (grant no. U24A2014, 32370852, 12090052, T2350007), the Ministry of Science and Technology of the People’s Republic of China (grant no. 2021ZD0201900, 2021YFA1101302).

## Competing interests

The authors declare no competing interests.

## Author information

### Contributions

J.S., X.C., F.C., H.C. and L.L. conceived the study and designed the experiments. J.S., L.M, X.L., H.S., L.L. and Z.W. supervised this study. J.S., L.M, X.L., L.L., Z.W., F.Y., G.Y., and J.S. contributed to resource acquisition. X.C., F.C., X.B., Y.C., H.S., H.C., Y.L. and J.J. performed biological and informatic experiments. X.C., J.L. designed and photographed the imaging characterization experiment. F.C., X.B., H.C., H.S., Y.L. and F.Y. analyzed the data. J.S., H.C., F.C., X.C. and J.X. wrote the paper. All authors read and provided feedback on the paper.

## Notes

### Competing Interest Statement

The authors have declared no competing interest.

### Summary of Updates

This version of the manuscript has been revised to update the author and affiliations.

