## Extended Data Fig. 1+ Extended Data Fig. 2 for "MAIS: an in-vitro sandbox enables adaptive neuromodulation via scalable neural interfaces"

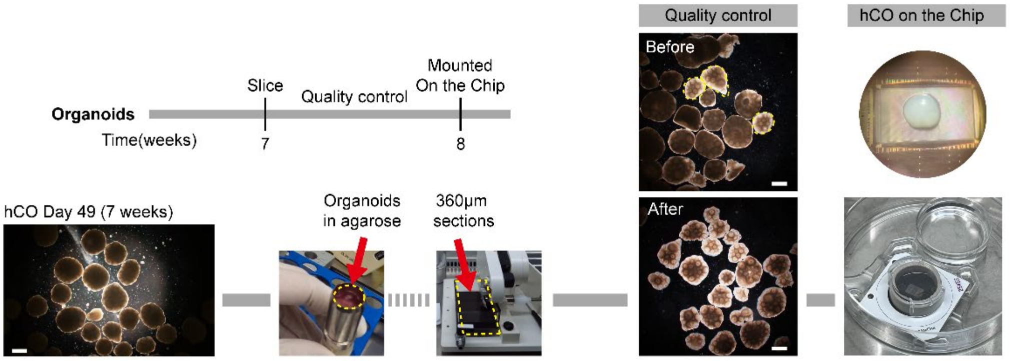


***Supplementary Fig. 1. Human brain organoid preparation for electrophysiology***
Cross-sectional slices were obtained from 7week-cultured hCOs. Whole organoids were embedded in 3% low-gelling-temperature agarose at ≈40 °C. Organoids embedded in the agarose gel were then sectioned into 360 μm thick slices with a vibratome. For recovery, slices were housed in transferred to ultralow-attachment plastic dishes. A recovery procedure of greater duration was designed, with a duration of one week, and a manual QC procedure was also added to the remove low-quality slices.

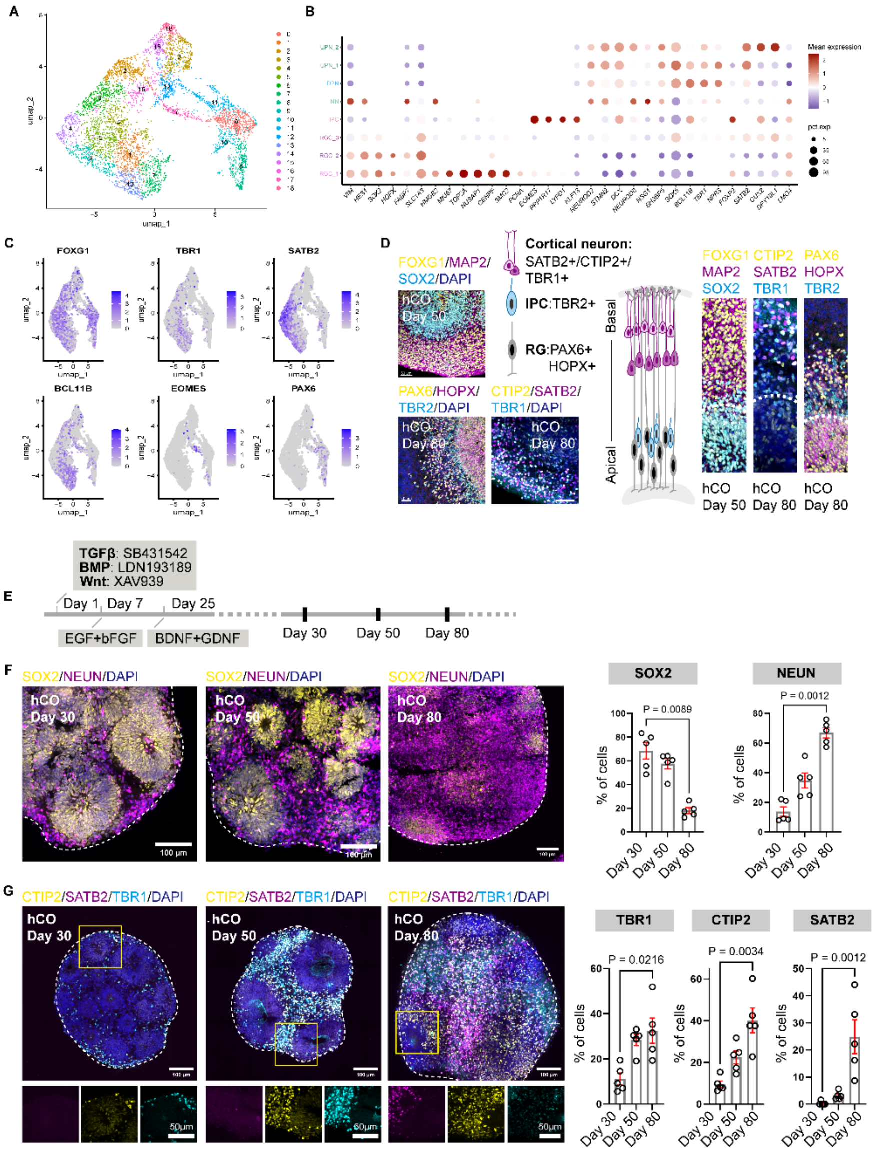


***Supplementary Fig. 2. Characterization of human cortical organoids in MAIS***

**a)** UMAP dimensionality reduction of snRNA-seq results from 80-day-old organoid cultures **b)** Annotating cell populations in an 80-day-old cortical organoid using specific markers. RGC: Radial glial cells; IPC: Intermediate progenitors; NN: Newborn neurons; DPN: Deep-layer neurons; UPN: Upper-layer neurons. **c)** snRNA-seq shows the expression of cortical-specific markers in the 80-day-old cortical organoid. **d)** Immunostaining for cortical-specific markers in Day 50/80 hCO; Progenitor markers (PAX6,HOPX,TBR2) and cortical neuron markers (TBR1,CTIP2,SATB2) show spatial distribution. Scale bar, 50 μm. **e)** Schematic of stained samples collection time. **f)** Immunostaining and quantifying SOX2/NEUN antibodies revealed neural differentiation in hCO. (n = 5 organoids). Scale bar, 100 μm. Data, mean ± SD. kruskal-wallis test. **g)** Immunostaining and quantifying CTIP2/TBR1/SATB2 antibodies revealed increased cortical neurons in hCO. (n = 5 organoids). Scale bar, 100 μm. Data, mean ± SD. kruskal-wallis test.
